# The physical state of the nucleus tracks its function across eukaryotes

**DOI:** 10.64898/2026.10.21.760156

**Authors:** Mirko Pleitner, Abin Biswas, Janek Uhlich, Simone Reber

**Affiliations:** Max Planck Institute for Infection Biology, Virchowweg 12, 10117 Berlin, Germany; Berliner Hochschule für Technik, Luxemburger Straße 10, 13353 Berlin, Germany; Max-Planck-Zentrum für Physik und Medizin, Kussmaulallee 2, 91054 Erlangen, Germany; Institute of Chemistry and Biochemistry, Freie Universität Berlin, 14195 Berlin, Germany

## Abstract

The nucleus is consistently less dense than the surrounding cytoplasm, a difference proposed to arise from selective nuclear import. The functional significance of low nuclear density, however, remains unknown. Because density is a bulk property of the cell, acute perturbations inevitably have pleiotropic consequences. We therefore turned to the natural experiments offered by the tree of life where nuclei vary in genome size, chromatin organisation, import selectivity and transcriptional state, and asked whether low nuclear density is tied to particular nuclear architectures or functions. Using dinoflagellates, Giardia and ciliates, we show nuclear density is uncoupled from genome size. In ciliates, two nuclei in one cytoplasm reach opposite densities relative to it, governed by nuclear import rather than transcription. We thus conform nuclear import as a major determinant of nuclear density and reveal nuclei that break the low-density rule.

## Introduction

The physical state of the nucleus shapes the reactions it hosts^1^, yet how it is set remains poorly understood. We recently showed that the nucleoplasm is consistently less dense than the surrounding cytoplasm across eukaryotic species, cell types and developmental stages, and that living systems actively maintain a constant nucleocytoplasmic density ratio^2^. We proposed that this is set by a pressure balance across the nuclear envelope, in which selective import establishes a nuclear proteome whose colloid osmotic pressure draws in water and reduces macromolecular density relative to the cytoplasm. The functional significance of low nuclear density, however, remains unknown. Because density is a bulk property of the cell, acute perturbations inevitably have pleiotropic consequences. We therefore turned to the natural experiments offered by the tree of life (Figure 1A, S1A), where nuclei vary in genome size, chromatin organisation, import selectivity and transcriptional state, and asked whether low nuclear density is tied to particular nuclear architectures or functions.

**Figure 1.**
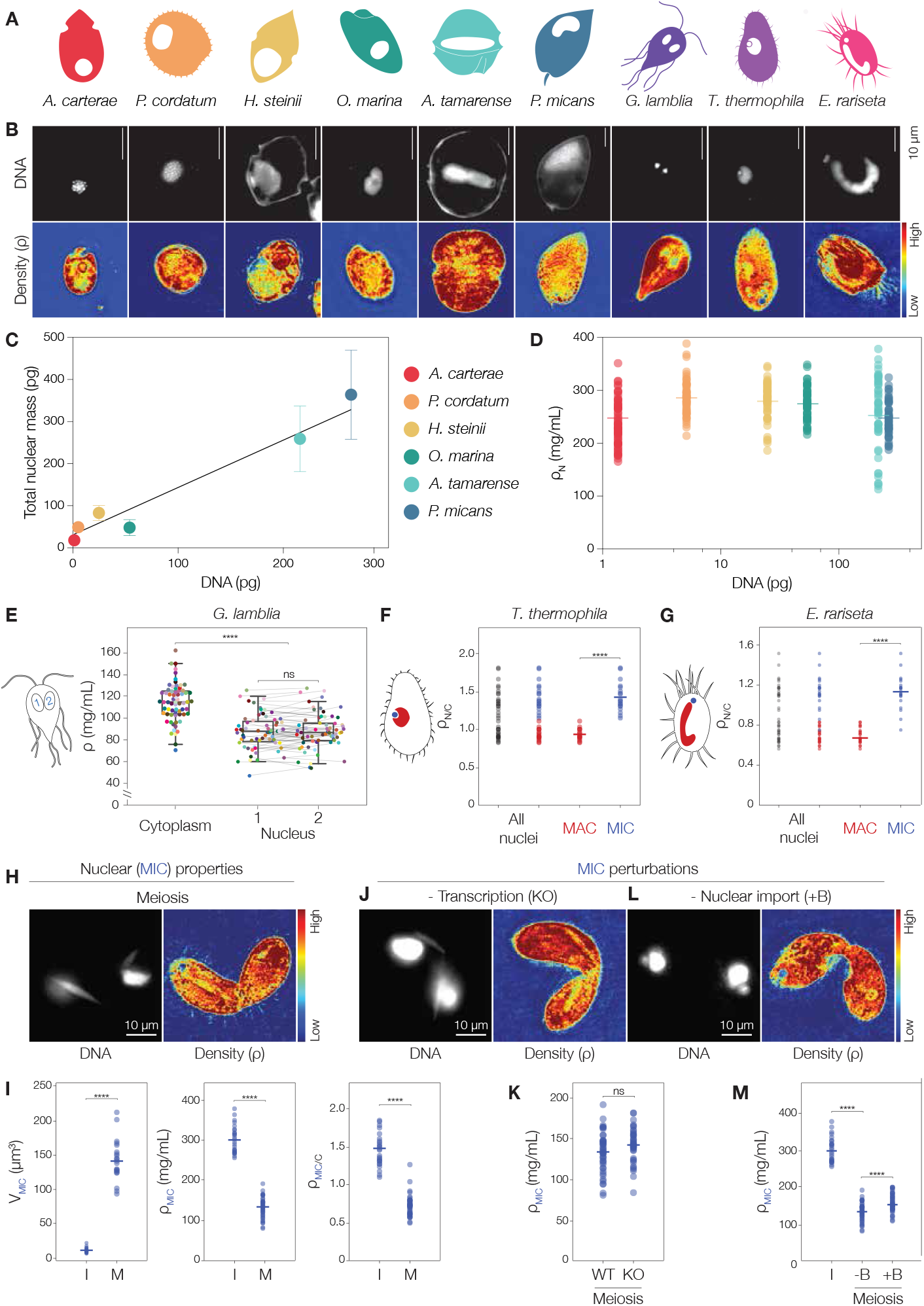
Statistical details can be found in Table S1. **(A)** Schematic representation of species with unconventional nuclear biology including Dinoflagellates (*A. carterae, P. cordatum, H. steinii, O. marina, A. tamarense, P. micans*), the common intestinal parasite *G. lamblia* and the ciliates *T. thermophila* and *E. rariseta*. Nuclear-to-cytoplasmic volume ratios drawn to scale. **(B)** Representative fluorescent images (DNA, upper panel) and central slice from reconstructed ODT tomograms showing density distributions from low to high (lower panel). **(C)** Total nuclear mass as measured by ODT versus DNA mass taken from published C-values (Table S1). Error bars SD. Weighted squares-linear regression with R^2^ = 0.725. **(D)** Nuclear density versus DNA mass. Bold line indicates the mean value. **(E)** Cytoplasmic and nuclear density in *G. lamblia*. Each data point represents the average mass density in either nucleus and the corresponding cytoplasm of a single cell. Grey lines connect two nuclei from the same cell indicated by identical colours. Mean±SD **(F)** Nuclear-to-cytoplasmic density ratio (ρ_N/C_). Mean nuclear density of either MAC (red) or MIC (blue) and the density of the corresponding cytoplasm in *T. thermophila*. **(G)** as in (F) for *E. rariseta*. **(H)** Representative fluorescent image (DNA) and central slice from reconstructed ODT tomograms showing density distributions of *T. thermophila* during prophase of meiosis. **(I)** *T. thermophila* MIC nuclear morphology in interphase (I) and meiosis (M). **(J)** Representative fluorescent image (DNA) and central slice from reconstructed ODT tomograms showing density distributions of *T. thermophila EMIT2* KO in meiosis. **(K)** Average MIC density in wildtype and *EMIT2* KO in meiosis. **(L)** Representative fluorescent image (DNA) and central slice from reconstructed ODT tomograms showing density distributions of *T. thermophila* nuclei in meiosis treated with benomyl (+B). **(M)** Average MIC density in interphase (I), meiosis (untreated, -B) and meiosis (benomyl-treated, +B). (E-G, I, K, M) Mann-Whitney U-test; ns indicates p > 0.05, **** indicates p < 0.0001.

## Results & Discussion

Dinoflagellates are among the most genomically extreme eukaryotes, largely nucleosome-free, maintaining hyper-condensed chromatin throughout the cell cycle, and possessing enormous genomes ranging up to 250 Gbp - over 80-fold the human haploid genome^3^. We imaged six species spanning essentially the full known range of dinoflagellate genome sizes from ~1–250 Gbp to identify nuclei and measure their mass density (ρ) using correlative fluorescence and optical diffraction tomography (ODT, Figure 1B, Table S1). Their extreme DNA content makes dinoflagellate nuclei dominated by DNA (Figure 1C), in contrast to most eukaryotic nuclei where DNA contributes at most 10% of total nuclear dry mass^2^. Across a 250-fold range in genome size, nuclear volume scaled linearly with total DNA mass and total nuclear dry mass (Figure S1D, E). Despite the large variation in genome content, nuclear density was maintained on average at 240–290 mg/mL across all six species (Figure 1D, S1F, G). Thus, across the full range of dinoflagellate genomes, nuclear volume scales with nuclear mass so that density remains constant, independent of genome size.

*Giardia lamblia* is a binucleate protist whose two nuclei share a common cytoplasm and are architecturally and functionally equivalent^4^. Both nuclei are diploid, contain complete copies of the genome, replicate simultaneously, and are transcriptionally active^5^. In our measurements, the two nuclei were less dense than the cytoplasm and indistinguishable in density (Figure 1E), consistent with their functional equivalence. Thus, two nuclei in one cytoplasm, equal in function, are equal in density. This raises the question of what happens when nuclear function differs.

Ciliates exhibit nuclear dimorphism: a single cell contains two morphologically and functionally distinct nuclei. The macronucleus (MAC) is the somatic nucleus, polyploid and transcriptionally active during vegetative growth. The diploid micronucleus (MIC) is the germline nucleus, transcriptionally inert except during meiosis^6^. We measured nuclear and cytoplasmic densities in two ciliate species, *Tetrahymena thermophila* and *Euplotes rariseta* (Figure 1F, G, S1H, I). In both, nucleocytoplasmic density ratios were broadly distributed, with some nuclei being less dense and some denser than the cytoplasm. Separating the data by nuclear identity revealed a striking dichotomy: in both species the MIC (blue) was consistently denser than the MAC (red). Notably, the MAC holds over an order of magnitude more DNA than the diploid MIC (~45n vs 2n), yet is the less dense of the two. This means that within a single cytoplasm, density can anti-correlate with DNA content. Together with the dinoflagellate data, this rules out DNA mass as the major determinant of nuclear density.

In *Tetrahymena*, the non-equivalent regulation of nuclear activities lies at least partly in the compositional difference of nuclear pore complexes and thus in transport selectivity^7^. The observed density divergence could therefore be explained by the MIC and MAC nuclear proteomes exerting distinct colloid osmotic pressures across the nuclear envelope. The only detectable MIC transcriptional activity occurs in meiosis^8^ enabled by the meiosis-specific import of Emit (Enhancer of meiotic transcription) proteins^9^. Meiotic prophase is accompanied by a well-documented elongation of the MIC^9^. Consistently, we found MIC volume to increase (Figure 1H,I). We additionally found that MIC density decreased, with the nucleocytoplasmic density ratio falling below 1 (Figure 1I, S1J). This dilution reflects the disproportionate growth of mass and volume: MIC dry mass increased 5.3-fold while volume increased 11.8-fold (Figure S1K). The MIC can thus reach a MAC-like low-density state when its nuclear biology changes. To distinguish transcriptional activity from nuclear import as the driver of the density reduction, we examined an *EMIT2* knockout in which the MIC remains transcriptionally silent during meiotic prophase^9^. MIC density was unchanged relative to wild type (Figure 1J, K), ruling out transcriptional activity as the driver of low nuclear density. While direct inhibition of nuclear import was unsuccessful, benomyl, which depolymerises microtubules, impaired MIC elongation (Figure 1L) as well as the meiotic gain in dry mass (Figure S1K), which led to a small but significant increase in MIC density during meiosis (Figure 1M). Together, these results are consistent with the idea that nuclear import establishes a specific nuclear proteome that draws water into the nucleus, reducing its macromolecular density below that of the cytoplasm. MIC density was not fully restored to interphase levels, most likely because benomyl blocks import only partially and indirectly, though we cannot exclude an additional import-independent contribution.

Our findings confirm that low nuclear density is the norm in *conventional*, import-competent nuclei, independent of genome size. Here, we identify nuclei that break this rule: nuclei that are denser than their cytoplasm (*P. micans*, interphase MIC in two Ciliates). Under the colloid-osmotic pressure model, a nucleus without protein import should relax toward cytoplasmic density^2^. Exceeding it requires an additional contribution from a component that is osmotically nearly inert but contributes a significant mass. Hypercondensed chromatin is one such case: it dominates mass while contributing minimally to osmotic pressure, the likely situation in *P. micans*, whose large genome is strongly condensed. A nuclear proteome enriched in large protein complexes would act similarly, adding mass on top of the osmotic pressure balance. For *P. micans*, these mechanisms remain speculative. For the *Tetrahymena* micronucleus, by contrast, we have direct evidence: its dense interphase state resolves during meiosis, and we show this transition is driven by nuclear import, not transcription.

## DECLARATION OF INTERESTS

The authors declare no competing interests.

## SUPPLEMENTAL INFORMATION

Supplemental information including methodological details and analyses and author contributions can be found with this article online.

## ACKNOWLEDGEMENTS

We thank Ricarda Leisering (Aebischer Group at the Robert Koch Institut Berlin) for *Giardia* strains, Gautam Dey (EMBL) for the *Euplotes* strain, Urban Tillmann (Alfred Wegener Institute) for *Oxyrrhis marina*, Kazufumi Mochizuki (IGH CNRS) for the *T. thermophila* strains and advice on *Tetrahymena* culturing and cell biology and Christian Goosmann (MPI for Infection Biology) for support at the Imaging Core Facility. We are grateful to Kyoohyun Kim (MPZ Erlangen) for optimising the script for field retrieval, tomogram reconstruction and axial drift correction. We thank all current and former members of the Reber lab, Vasily Zaburdaev, Iain Patten and Alf Herzig for critical discussions. The authors acknowledge funding from the DFG (528483508 - FIP 12 to SR) and the Max Planck Society (to AB, SR).

## Supplemental Information

**Supplemental Figure S1.**
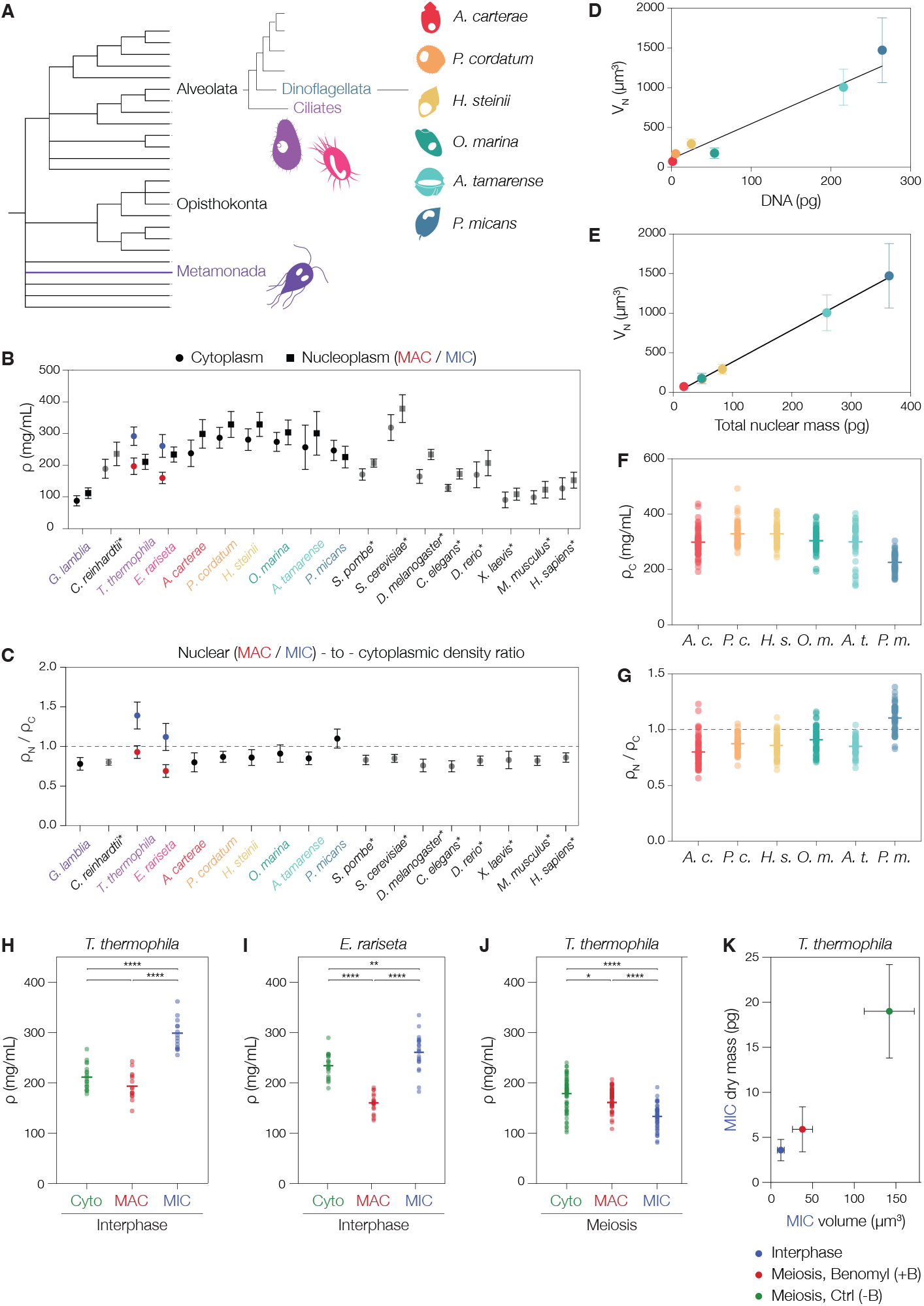
Statistical details can be found in Table S1. **(A)** Phylogeny based on^S1,S2^. **(B)** Absolute values of nucleoplasmic densities (circles) and cytoplasmic densities (squares) from 18 eukaryotes. For *T. thermophila* and *E. rariseta*, the densities of the MAC (red) and MIC (blue) have been plotted separately. Error bars SD. Values in grey and species indicated with an asterisk indicate data previously published in^S3^. Black dashed line: N/C density = 1. **(C)** N/C density ratios from 18 eukaryotes. **(D)** Each data point represents the average nuclear volume and the DNA mass taken from published C-values. Error bars SD. Weighted squares-linear regression with R^2^ = 0.770. **(E)** Each data point represents the average nuclear volume and the average total nuclear mass. Error bars SD. Linear regression with R^2^ = 0.998 **(F)** Mean cytoplasmic density. Each data point represents the value from the cytoplasm of a single cell. The bold line indicates the mean value. **(G)** N/C density ratio. Each data point represents the value from a single cell. The bold line indicates the mean value, the dashed line visualises an N/C density ratio of 1. **(H)** Dry mass density of interphase cytoplasm (green), MAC (red) and MIC (blue) in *T. thermophila*. **(I)** Dry mass density of the interphase cytoplasm (green), MAC (red) and MIC (blue) in *E. rariseta*. **(J)** Dry mass density of the cytoplasm (green), MAC (red) and MIC (blue) during meiotic prophase of *T. thermophila*. (H-J) Mann-Whitney, ns indicates p > 0.05, * indicates p < 0.05, ** indicates p < 0.01 and **** indicates p < 0.0001. **(K)** MIC dry mass versus volume in *T. thermophila* during interphase (blue, I), meiotic prophase untreated (green, -B) and meiotic prophase treated with Benomyl (red, +B).

**Supplemental Table S1:** DNA content, nuclear volume and mass of *A. carterae, P. cordatum, H. steinii, O. marina, A. tamarense, P. micans, G. lamblia, T. thermophile* and E. *rariseta*.

| Data from literature |  |  |  | Condition | Fluorescence volume data | ODT measurements |  |  |  |  |  |
| --- | --- | --- | --- | --- | --- | --- | --- | --- | --- | --- | --- |
| Species | Genome | DNA (pg) | Reference |  | n nuclei | n nuclei | N/C ratio | ρ [mg/ml] μ ± SD | ROIs | i.i.d. | Statistical test |
| *Amphidinium carterae* | 1.25 Gbp | 1.35 | Judd et al. 2025 | interphase | 22 | 71 | 0.80 ± 0.12 | ρ<sub>N</sub>=238±42<br>ρ<sub>C</sub>=299±45 | 10 ROIs | 4 | Weighted least squares-linear regression of pooled biol. replicates: total mass vs DNA mass, density vs. DNA mass |
| *Prorocentrum cordatum* | 4.15 Gbp | 4.48 | Kalvelage et al. 2023 | interphase | 20 | 61 | 0.87 ± 0.07 | ρ<sub>N</sub>=287±33<br>ρ<sub>C</sub>=329±41 | 10 ROIs | 3 |  |
|  | 4.75 Gbp | 5.13 | Dougan et al. 2023 |  |  |  |  |  |  |  |  |
| *Heterocapsa steinii (triquetra)* | 23 Gbp | 24.83 | Mc Ewan et al. 2008 | interphase | 22 | 60 | 0.86 ± 0.10 | ρ<sub>N</sub>=281±30<br>ρ<sub>C</sub>=329±38 | 10 ROIs | 3 |  |
| *Oxyrrhis marina* | 50 Gbp | 53.97 | Sano and Kato, 2009 | interphase | 23 | 60 | 0.91± 0.11 | ρ<sub>N</sub>=274±30<br>ρ<sub>C</sub>=304±39 | 10 ROIs | 5 |  |
| *Alexandrium tamarense* | 200 Gbp | 215.87 | Wang et al. 2024 | interphase | 19 | 49 | 0.85 ± 0.08 | ρ<sub>N</sub>=257±70<br>ρ<sub>C</sub>=301±69 | 10 ROIs | 3 |  |
| *Prorocentrum micans* | 245 Gbp | 264.45 | Hou & Lin 2009 | interphase | 17 | 53 | 1.10 ± 0.12 | ρ<sub>N</sub>=247±32<br>ρ<sub>C</sub>=226±31 | 10 ROIs | 6 |  |
| *Giardia lamblia* | 11.7 Mbp (1 N) | 0.0252 / nucleus (2N) | Xu et al. 2020 | interphase | - | nucleu s 1: 67<br>nucleu s 2: 67 | 0.78 ± 0.08<br>0.79± 0.08 | ρ<sub>N_avg</sub>=88.5±16<br>ρ<sub>N1</sub>=88±16<br>ρ<sub>N2</sub>=89±16<br>ρ<sub>C</sub> =112±17 | 5 ROIs per nucleus | 5 | Mann-Whitney U test of pooled biol. replicates: cytoplasm vs. combined nuclei and nucleus 1 vs nucleus 2 |
| *T. thermophila* - MAC | 103.34 Mbp (1N) + 21kb rDNA minichromosome, ~9000 copies | 5.02 / nucleus (45N)<br>5.22 including rDNA | Ye et al. 2025, Sheng et al. 2020 | interphase | 20 | 17 | 0.93 ± 0.07 | ρ<sub>MAC</sub>=197±26<br>ρ<sub>Cyto</sub> =211±24 | 10 ROIs | 3 | Mann-Whitney U test of pooled biol. replicates: MAC vs MIC<br>In Fig.S1: Cyto vs. MAC, Cyto vs. MIC |
|  |  |  |  | Meiosis | 21 | 49 | 0.96± 0.16 | ρ<sub>MAC</sub> = 168±23<br>ρ<sub>Cyto</sub> = 179±34 | 10 ROIs | 3 | Mann-Whitney U test of pooled biol. replicates: MAC vs MIC<br>In Fig.S1: Cyto vs. MAC, Cyto vs. MIC |
|  |  |  |  | *EMIT2*-KO | - | 43 | - | - | 10 ROIs | 3 | - |
|  |  |  |  | Benomyl | 20 | 47 | 0.97± 0.10 | - | 10 ROIs | 4 | - |
| *T. thermophila* - MIC | 157 Mbp (1N) | 0.34 / nucleus (2N) | Hamilton et al. 2016 | interphase | 20 | 17 | 1.39 ± 0.17 | ρ<sub>MIC</sub>= 292±29<br>ρ<sub>Cyto</sub> = 211±24 | 1 ROI | 3 | Mann-Whitney U test of pooled biol. replicates: MAC vs MIC |
|  |  |  |  |  |  |  |  |  |  |  | In Fig.S1: Cyto vs. MAC, Cyto vs. MIC |
| | | | | Meiosis | 21 | 45 | $0.79 \pm 0.12$ | $\rho_{\text{MIC}} = 133 \pm 24$<br>$\rho_{\text{Cyto}} = 179 \pm 34$ | 10 ROIs | 3 | Mann-Whitney U test of pooled biol. replicates: MIC_interphase vs. MIC_meiosis<br>In Fig.S1: Cyto vs. MAC, Cyto vs. MIC |
| | | | | EMIT2-KO | - | 43 | - | $\rho_{\text{MIC}} = 142 \pm 20$ | 10 ROIs | 3 | Mann-Whitney U test of pooled biol. replicates: MIC_WT_meiosis vs. MIC_EMIT2KO_meiosis |
| | | | | Benomyl | 20 | 47 | $0.81 \pm 0.08$ | $\rho_{\text{MIC}} = 157 \pm 23$ | 2-10 ROIs | 4 | Mann-Whitney U test of pooled biol. replicates: MIC_interphase vs. MIC_meiosis_untreated vs. MIC_meiosis_benomyl |
| *E. rariseta* -<br>MAC | N.d. | | | interphase | - | 21 | $0.69 \pm 0.08$ | $\rho_{\text{MAC}} = 160 \pm 18$ | 10 ROIs | 3 | Mann-Whitney U test of pooled biol. replicates: MAC vs MIC<br>In Fig.S1: Cyto vs. MAC, Cyto vs. MIC |
| *E. rariseta* -<br>MIC | N.d. | | | interphase | - | 21 | $1.12 \pm 0.18$ | $\rho_{\text{MIC}} = 261 \pm 36$ | 1 ROI | 3 | Mann-Whitney U test of pooled biol. replicates: MAC vs MIC<br>In Fig.S1: Cyto vs. MAC, Cyto vs. MIC |

Convert DNA length to mass:
Average molecular weight of a double-stranded DNA base pair: 650 Dalton (g/mol)

## Methods

### Strains

*A. carterae*: RCC1522 - https://roscoff-culture-collection.org/rcc-strain-details/1522

*P. cordatum*: RCC291 - https://roscoff-culture-collection.org/rcc-strain-details/291

*H. steinii:* RCC2540 - https://roscoff-culture-collection.org/rcc-strain-details/2540

*O. marina*: a gift from Urban Tillmann, AWI

*A. tamarense*: RCC292 - https://roscoff-culture-collection.org/rcc-strain-details/292

*P. micans*: RCC3046 - https://roscoff-culture-collection.org/rcc-strain-details/3046

*G. lamblia*: WB-C6, ATCC 50803

*T. thermophila*: *B2086* and *CU428, EMIT2-KO*: *EMIT2-KO-S5-1* and *EMIT2-KO-S9-3*

*E. rariseta*: a gift from Gautam Day, EMBL

### DNA content

The DNA content per nucleus was taken from published C-values (Table S1).

### Correlative confocal fluorescence and optical diffraction tomography (ODT)

A custom-built correlative confocal fluorescence and ODT setup was used in this study. Further details on the setup, image acquisition and tomogram reconstruction can be found in ^S3,S4,S5^. Briefly, for imaging, live cells were stained with HOECHST 33342 and transferred onto imaging dishes. The holograms were recorded and reconstructed and each voxel within the 3D RI tomogram was assigned a refractive index (RI) value. To obtain RI information, square ROIs with sizes of 1×1 μm and 1.5×1.5 μm were manually and randomly placed in the central xy slice of the tomogram within nuclei and the cytoplasm. In the cytoplasm, clearly distinguishable organelles such as vacuoles, chloroplasts, starch granules and pigment bodies were avoided for quantification. Dry mass density was calculated using the RI of the medium, measured with an Abbe refractometer, and the RI α-increment of proteins and nucleic acids.

### Giardia

In Figure 1E, each data point represents the average mass density of 5 randomly distributed ROIs in either nucleus and 10 randomly distributed ROIs the corresponding cytoplasm of a single cell. We failed to detect a statistically significant within-cell difference of both nuclei (paired t-test, p =0.1253) and can exclude inter-nuclear differences greater than 3.19 mg/mL (90% CI), corresponding to 0.2*SD of nuclear mass density.

### Tetrahymena

Conjugation was initiated by mixing equal volumes of starved wild type B2086 and CU428 strains or starved *EMIT2-KO-S5-1* and *EMIT2-KO-S9-3* strains. In Figure 1K each data point represents the average of 10 randomly distributed ROIs per MIC in the wild type or the KO during meiosis. In Figure 1M, each data point represents the average of 10 randomly distributed ROIs in interphase and 2 randomly distributed ROIs in meiosis. For benomyl treatment, a final concentration of 20 µg/ml was used.

### Declaration of generative AI and AI-assisted technologies in the manuscript preparation process

During the preparation of this work, the author(s) used Claude by Anthropic to refine the text for conciseness and to meet word count requirements. The author(s) reviewed and edited the output and take full responsibility for the content of the published article.

## Author contributions

MP cell culture, image and data analysis, AB tomogram reconstruction, image and data analysis, JU perturbation experiments, image and data analysis, SR conceived and supervised the project and wrote the paper with input from all authors.

## Notes

### Competing Interest Statement

The authors have declared no competing interest.

